# Mitochondrial genome reduction and accelerated evolution in planktonic foraminiferans

**DOI:** 10.1101/2025.08.28.672841

**Authors:** Hsin-Tung Lai, Ming-Wei Lai, Tzu-Haw Wang, Haojia Ren, Chuan Ku

**Affiliations:** Institute of Plant and Microbial Biology, Academia Sinica, Taipei, Taiwan; Institute of Ecology and Evolutionary Biology, National Taiwan University, Taipei, Taiwan; Department of Geosciences, National Taiwan University, Taipei, Taiwan

**Keywords:** Comparative genomics, Foraminifera, mitogenome, protist, reductive evolution, Rhizaria, single-cell genomics

## Abstract

The evolution of mitochondria provides crucial insights into the diversification of eukaryotes, with complex events of gene losses revealed through comparative analyses of mitochondrial genomes (mitogenomes) across eukaryotic lineages. However, the mitogenomes of many microbial eukaryotes remain underexplored due to challenges in their isolation and cultivation. Particularly understudied are Foraminifera (Rhizaria, SAR), unicellular calcifyers that are widely distributed across global oceans and important paleoenvironmental proxies. Through single-cell genomic sequencing, here we report a 22-kb complete mitogenome of a planktonic foraminiferan from tropical seawater, which is the smallest known to date among all sequenced mitogenomes of Rhizaria, a major lineage of eukaryotes. It contains only six protein-coding genes and fragmented ribosomal RNA genes, and has lost most genes in oxidative phosphorylation and all genes in mitochondrial translation. Such genome reduction is associated with accelerated evolutionary rates and a lower GC content than that of benthic foraminiferan and other rhizarian mitogenomes. These findings highlight the unique trajectory of mitogenome reduction during rhizarian evolution and the use of single-cell approaches for recovering microbial eukaryotic genomes and expanding our understanding of mitochondrial evolution.

## Introduction

Mitochondria are endosymbiotic organelles that originated from free-living alphaproteobacteria [1,2]. Over time, mitochondrial genomes have undergone gene loss and endosymbiotic gene transfer to the nuclear genome, resulting in extensive gene reduction and diversification of mitogenome functions and structures [3-6]. Animals and plants have more than 20,000 complete mitogenome sequences in the NCBI Organelle Database (released on Feb. 12, 2025). In contrast, many microbial eukaryote taxa, particularly those that cannot be easily cultured, have few mitogenome sequence records [7], although their mitogenomes demonstrate greater diversity [8]. It is thus necessary to have more extensive sampling of microbial eukaryotes for better understanding of mitogenome evolution.

As the primary site of energy production, mitochondria typically retain genes encoding proteins involved in oxidative phosphorylation (electron transport chain [ETC; complexes I-IV] and ATP synthase [complex V]) and translation (ribosomal proteins, rRNA, and tRNA) [9]. Mitogenomes show remarkable variation in gene contents, with up to 66 protein-coding genes in *Andalucia godoyi* (Discoba) [10]. Different degrees of mitogenome reduction are observed across the tree of eukaryotes, with complete loss of mitochondrial DNA in diverse anaerobes that have the derived forms of mitochondria, hydrogenosomes or mitosomes [11]. In the facultative anaerobic alga *Euglena gracilis* (Discoba), which can use fumarate as terminal electron acceptors during anoxia, the mitogenome has only 5 protein-coding genes and its rRNA genes are fragmented [11,12]. In diverse myzozoans (Alveolata), including apicomplexan parasites, dinoflagellates, and *Chromera*, mitochondria generally have fewer than 5 protein-coding genes and their ribosomes contain dozens of small rRNA fragments [13-15].

Rhizaria, which comprises Cercozoa, Endomyxa, and Retaria [16], are diverse free-living or parasitic, amoeboid eukaryotes and one of the least studied eukaryotic supergroups in terms of mitogenomes, with only 12 reference mitogenome sequences (4 cercozoans, 4 endomyxea, 4 retarians) currently available. In Retaria, Foraminifera are marine, mostly calcifying, large unicellular eukaryotes, living either as free-floating planktonic or bottom-dwelling benthic organisms. Due to their wide distribution and preservable calcareous shells, foraminiferans play crucial roles in carbon cycling and serve as important bioindicators and microfossils in paleoceanographic studies [17,18].

Available genomic data of Foraminifera are mostly limited to specific gene fragments (e.g. nuclear SSU rDNA [19,20], mitochondrial *COX1* gene [21]) obtained from environmental samples. Whole-genome sequencing data exist for only three benthic species, *Astrammina rara* [22], *Globobulimina* sp. [23], and *Reticulomyxa filosa* [24]. In addition, circularized mitogenome sequences have been reported from two benthic foraminiferans, *Neorotalia gaimardi* and *Calcarina hispida*, which have reduced gene contents that lack all of the ribosomal protein genes [25]. Until to date, no whole-genome sequencing data are currently available for planktonic foraminifera, despite their ecological significance and global fossil records.

In this study, we applied a single-cell (single-particle) approach to genome sequencing for planktonic foraminifera. This culture-independent method enables the examination of sequence information from individual cells and has important potential for the study of microbial eukaryotes [26-29]. Although the coverage of nuclear genomes has been limited by the low-copy nature of the nuclear DNA molecules, single-cell genomics can generate complete genome sequences of the mitochondria [25,30-31], which can have hundreds to thousands of copies within a cell [32]. Using this approach, we recovered a complete, circularized mitogenome sequence of a planktonic foraminiferan isolated from tropical seawater and conducted comparative and phylogenomic analyses of foraminiferan mitogenomes.

## Materials and Methods

### Sample collection

Marine particles were collected in August 2023 by towing a 200-µm-mesh plankton net at a depth of ∼13 meters near Gong-Guan Harbor, Green Island, Taitung County, Taiwan (22.69°N, 121.48°E), where planktonic foraminiferans have been observed [33,34]. COPAS Vision, a large-particle imaging flow cytometer (Union Biometrica, Holliston, USA), was used for fast preliminary examination of particle composition. To preserve the intactness of foraminiferan particles, we carefully isolated individual particles manually with a micropipette and observed them under a Nikon ECLIPSE Ts2-FL inverted optical microscope (Nikon, Tokyo, Japan). The particles were washed with artificial seawater, individually transferred into a 96-well plate with each well containing 100 µL 5 % DMSO in artificial seawater, and preserved in -20 °C until further processing for DNA extraction.

### Sample preparation for SEM observation

For SEM observation, 40 mL of the net tow material was fixed with 1% formaldehyde and 0.05% glutaraldehyde to preserve microbial morphology. The sample was filtered onto a 0.2 µm PC membrane and rinsed with phosphate buffer, followed by ethanol dehydration and critical point drying (EM CPD300: Leica, Wetzlar, Germany). Finally, it was cut into sections and mounted onto stubs for SEM imaging (FEI Quanta 200: FEI, Hillsboro, USA).

### DNA extraction and sequencing

Single foraminiferan particles were individually transferred to 1.5-mL Eppendorf tubes and ground into powder using a customized plastic stick with a granular head that fits the bottom of the Eppendorf tube. Cell lysis and whole-genome amplification were carried out using the REPLI-g Advanced DNA Single Cell Kit (QIAGEN, Hilden, Germany) following the manufacturer’s protocol. The amplified DNA fragments were purified using the AMPure XP Bead-Based Reagent (Beckman Coulter Life Science, Brea, USA). Purified DNA samples were quantified using Qubit HS (Thermo Fisher Scientific, Waltham, USA). Libraries were prepared using Illumina DNA Prep (Illumina, San Diego, USA), with the lengths estimated by Fragment Analyzer (Agilent, Santa Clara, USA), and sequenced on the Illumina NovaSeq 6000 platform (150-bp paired-end reads). The remaining DNA samples of three particles (GI-A, GI-C, and GI-D) were also sequenced using Nanopore R10.4.1 flowcell (Oxford Nanopore Technologies, Oxford, UK).

### Genome assembly

Illumina short reads were processed with fastp v0.22.0 [35] for adapter removal and trimming. For the Nanopore long reads, real-time basecalling was performed using MinION MK1C (Oxford Nanopore Technologies, Oxford), and the reads were trimmed using Porechop v0.2.4 [36]. Illumina short reads were assembled using SPAdes v4.0.0 [37] in single-cell mode (--sc) with specified k-mers (-k 21, 33,55,77,99,127). Assembled contigs were then classified by Tiara v1.0.3 [38] into 6 groups based on k-mer frequencies (Archaea, Bacteria, Prokarya, Eukarya, Plastid, Mitochondrion). The contigs assigned to the group “Mitochondrion” were searched using the *blastx* command in BLAST+ v2.13.0 [39] against a custom protein database containing protein-coding sequences of the two available foraminiferan mitogenomes (benthic species *Calcarina hispida* [OP965950] and *Neorotalia gaimardi* [OP965949] [25]) and representatives of Radiolaria (Acanthometra sp. and Lithomelissa sp. [25]) and Cercozoa (*Lotharella oceanica* [NC_029731] [40]) using an E-value cutoff of 1e-10. Mitochondrial contigs with hits to this database were extracted and individually served as seed inputs to NOVOPlasty v4.3.1 [41].

### Quality assessment of genome assembly

Illumina and Nanopore reads were mapped to the best assembled mitogenome sequence (Globigerinidae sp. GI-D) using Bowtie2 v2.4.1 [42] and Minimap2 v2.28-r1209 [43], respectively, for visual inspection in Integrative Genomics Viewer (IGV) [44] to ensure the correctness of the genome assembly. The circularity and completeness of the mitogenome were confirmed by examining the contigutiy of both long and short reads and by shifting the endpoints of the assembly by 1 kb. The quality of the circularized mitogenome sequences was further evaluated by QUAST v5.0.2 [45]. The repetitive sequence analysis in the mitogenome was conducted by PERF v0.4.6 [46].

### Genome annotation

The Globigerinidae sp. GI-D mitogenome was annotaed using the MFannot web server [47] and MITOS2 [48] with reference genome set refseq63o. The genetic code was set to NCBI genetic code 4 (the mold, protozoan, and coelenterate mitochondrial code, and the mycoplasma/spiroplasma code) for annotating mitochondrial protein-coding, tRNA, and rRNA genes. Unannotated regions were translated into amino acid sequences also using the NCBI genetic code 4 and searched against the predicted protein sequences from the two available foraminiferan mitogenomes (*Calcarina hispida* and *Neorotalia gaimardi* [25]) using *blastx*. The annotations were visualized using the OGDRAW web server [49].

### Phylogenetic analyses

Amino acid sequences of whole mitogenome protein-coding genes of Globigerinidae sp. GI-D and other Rhizaria species, retrieved from NCBI and previous publications (**Supplementary Table 1**), were aligned and concatenated using Orthofinder v2.5.5 [50]. A maximum likelihood phylogeny was inferred with IQ-TREE v2.2.2.7 [51], using ModelFinder [52] to determine the best-fit substitution model according to the Bayesian information criterion (BIC). The final model was selected manually, taking into account the intended use and assumptions underlying each model. For individual gene trees, sequence alignments generated with MAFFT v7.520 [53] and phylogenies were reconstructed using IQ-TREE in the same way. Phylogenetic trees were visualized in FigTree v1.4.4 [54].

## Results

### Complete mitogenome sequence from a planktonic foraminiferan

In total, 20 foraminiferan particles from the family Globigerinidae (Rotaliida, Globothalamea) were isolated, which was the dominant taxon of planktonic foraminiferans in seawater offshore Green Island, Taiwan. After single-cell genome amplification, we obtained enough DNA for four particles and sequenced their DNA. Among them, the single amplified genome (SAG) from the particle Globigerinidae sp. GI-D has the best assembly quality supported by both Illumina and Nanopore reads (**Table 1**). Although each particle was thoroughly washed with aseptic artificial seawater, putative prokaryotic contigs that likely came from closely associated or attached prokaryotic cells still comprised the majority of all contigs for each particle, with GI-D having the highest proportion of eukaryotic contigs (**Supplementary Table 2**). To recover the authentic Foraminifera genomes, only eukaryotic contigs identified by Tiara were retained for further analyses. Contigs showing high similarity to mitochondrial protein-coding sequences from the two benthic foraminiferans, *N. gaimardi* and *C. hispida*, were identified and used as seed sequences (baits) to assemble more complete mitochondrial genomes using NOVOPlasty. This approach successfully recovered a circularized mitogenome sequence of the particle GI-D (GenBank accession PV558267), which is 22,652 bp in length and has a GC content of 21.5% (**Figure 1**). Mapping of both Illumina short reads (7435x coverage) and Nanopore long reads (560x coverage) confirmed the contiguity of read coverage and the correctness of the assembly, including the circularity of the mitogenome. In addition, a 30,272-bp circularized mitogenome sequence was recovered from a *Chrysochromulina* sp. (Prymnesiales, Haptista) associated with GI-D (GenBank accession PV535690). Partial mitogenome sequences were assembled from two other planktonic foraminiferan particles, GI-A (2,279 bp) and GI-C (6,068 bp), whose mitochondrial *cox1* sequences are closely related to that of GI-D (**Supplementary Figure 1**).

**Table 1.** Assembly statistics of the Globigerinidae sp. GI-D SAG

|  | **Globigerinidae sp. GI-D SAG** |
| --- | --- |
| Illumina read data | 20828.54 Mb |
| Nanopore read data | 1464.86 Mb |
| # assembled contigs > 1000 bp | 30,626 |
| Assembly size | 180.6 Mb |
| GC content | 43.96 % |
| N50 | 9,014 |
| L50 | 3,102 |
| Illumina read coverage | 153x |

**Figure 1.**
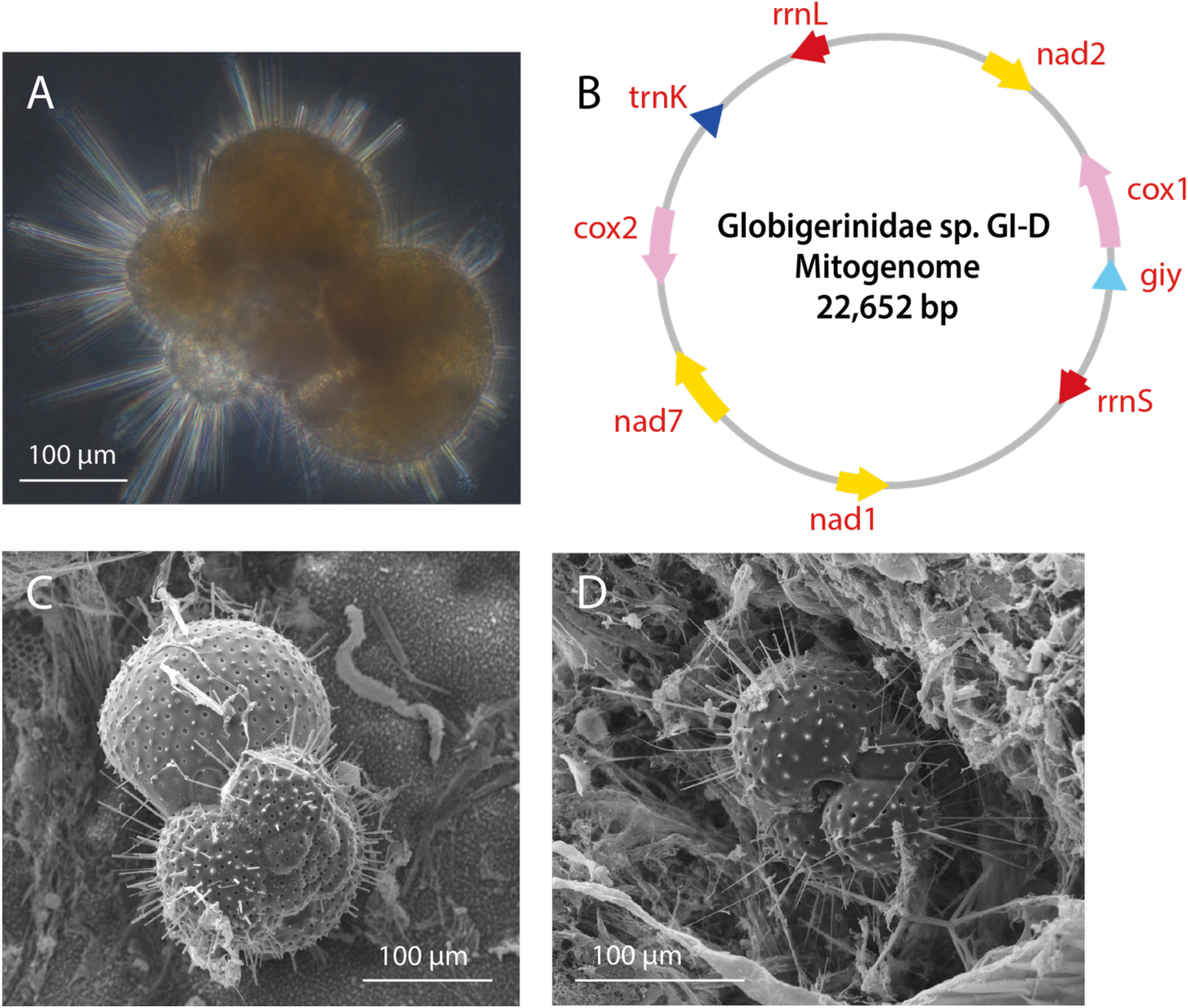
Globigerinidae and its mitochondrial genome sequence. (A) The planktonic foraminiferan particle GI-D from the family Globigerinidae before DNA extraction. (B) Mitogenome map of GI-D, showing genes in diflerent functional categories: yellow (complex I), pink (complex IV), red (ribosomal RNA), dark blue (tRNA), and light blue (others). Arrows indicate the transcriptional direction of each gene. (C-D) SEM images of morphologically similar particles from Globigerinidae in the same seawater sample.

The particle GI-D is multilocular with distinct spines extending from the chamber walls (**Figure 1**). Its globular chambers are arranged in a trochospiral coiling pattern, with 4 chambers clearly visible in the final whorl and smaller chambers coiled centrally. These morphological features strongly suggest that GI-D is affiliated to the family Globigerinidae, as Globigerinidae spp. are spinose and exhibit trochospiral or near-planispiral chamber arrangements [55]. The chamber wall surface of GI-D is rough and perforated, with yellow-green color, which is likely due to symbiotic microalgae. Because only the lateral view of GI-D was photographed, the aperture (the opening to exterior on the final chamber) was unobservable, making it difficult to classify GI-D to the species level.

### Substantial mitogenome reduction and gene losses

The mitogenome of Globigerinidae sp. GI-D is by far the smallest mitogenome among all rhizarian mitogenomes and contains only 6 protein-coding genes and a single tRNA gene (trnK) (**Figure 1**). Similar to other retarians, the GI-D mitogenome only has fragmented large- and small-subunit rRNA genes, in contrast to the full-length rRNA genes found in cercozoans, stramenopiles, and alveolates (**Figure 2**). The retarian (Foraminifera and Radiolaria) mitogenomes have substantially reduced gene contents, characterized by the loss of *nad6, nad9, atp8*, and all ribosomal protein genes (**Figure 2**). The proportion of gene-coding regions in Foraminifera (17.4∼33.5%) is significantly lower than in other rhizarian mitogenomes (e.g. *Lotharella oceanica*: 95%; *Spongospora subterranea*: 66%), and the human mitogenome (93% [56]) (**Table 2**).

**Table 2.** Mitogenome information of foraminiferan species.

|  | Mitogenome size | GC content | Gene-coding region | Repetitive content | Repeat frequency/kb | Accession ID |
| --- | --- | --- | --- | --- | --- | --- |
| GI-D | 22, 652 bp | 21.5% | 23.3% | 3.6% | 3.62 | PV558267 |
| *C. hispida* | 46,016 bp | 23.5% | 17.4% | 1.4% | 1.15 | OP965950 |
| *N. gaimardi* | 50,389 bp | 22.7% | 33.5% | 1.1% | 1.49 | OP965949 |

**Figure 2.**
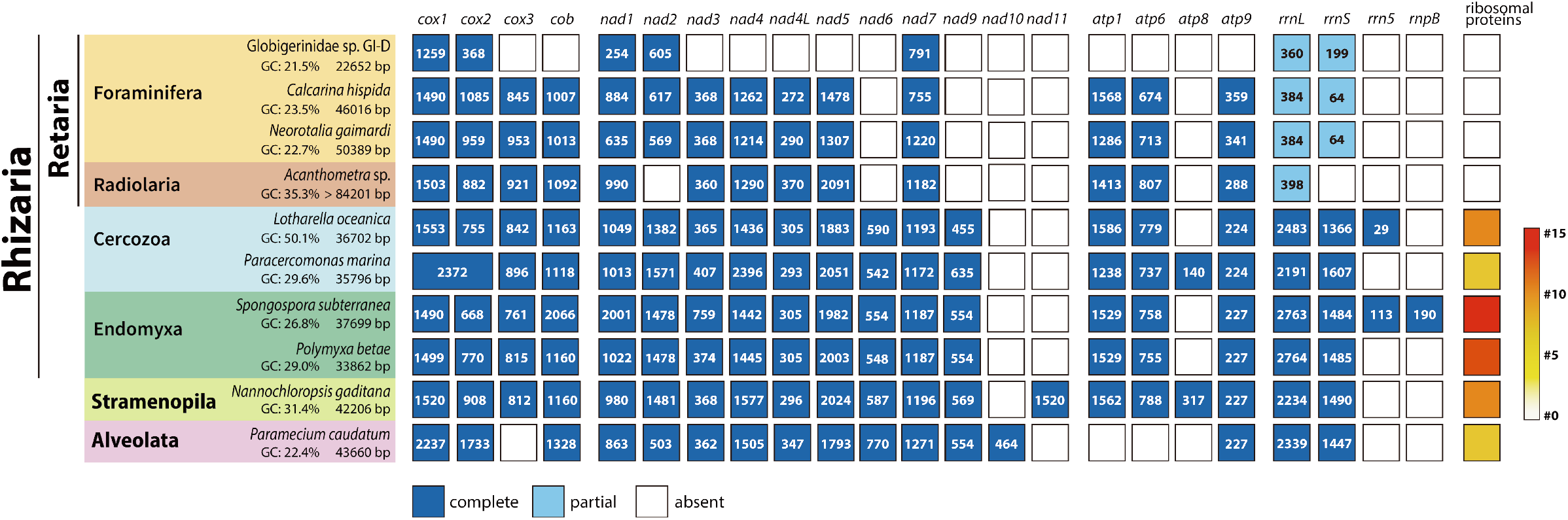
Comparative mitochondrial genomics of Rhizaria. The presence, partial presence, and absence of genes in mitogenomes from Rhizaria and representatives of Stramenopila and Alveolata are indicated by dark blue, light blue, and white blocks, respectively. Numbers within dark or light blue blocks represent gene lengths in bp. For ribosomal proteins, a red-to-white gradient indicates the number of ribosomal proteins encoded in each mitogenome.

The planktonic foraminiferan GI-D has a mitogenome that is even more reduced in both genome size and gene content compared with the two benthic foraminiferans *Calcarina hispida* and *Neorotalia gaimardi*. Specifically, *cox3, cob, nad3, nad4, nad4L, nad5*, and all *atp* subunit genes are absent. Although small fragments with similarity to *cox3, nad5*, and *atp1* could be detected through MITOS2, these fragments are all less than 50 bp. On the other hand, the Globigerinidae sp. GI-D mitogenome is uniquely annotated with a partial *giy* gene (encoding GIY-YIG protein) and *trnK*, which are absent in the benthic species. Additionally, GI-D has a lower GC content and a higher proportion of repetitive sequences in the non-coding regions (**Table 2** and **Supplementary Figure 2**).

### Phylogenetics and accelerated evolution of foraminiferan mitogenomes

A mitogenome tree of Rhizaria was reconstructed from whole-mitogenome protein sequences (**Figure 3**), showing the grouping of Globigerinidae sp. GI-D with the two benthic foraminiferans *Calcarina hispida* and *Neorotalia gaimardi*. To expand the taxon sampling of foraminiferans to beyond those with whole-mitogenome sequences, another tree was built from the mitochondrial COX1 sequence, which has a good balance of conservation and variation for resolving the relationships between foraminiferans [57]. The COX1 tree suggests that GI-D is sister to *Orbulina universa* [58], which is a known member of the family Globigerinidae [55] (**Figure 4**). The comparative genomic (**Figure 2**) and phylogenetic (**Figure 3** and **Figure 4**) analyses of mitogenomes also suggest that the two benthic species, which both belong to the family Calcarinidae, are closer to each other than to GI-D.

**Figure 3.**
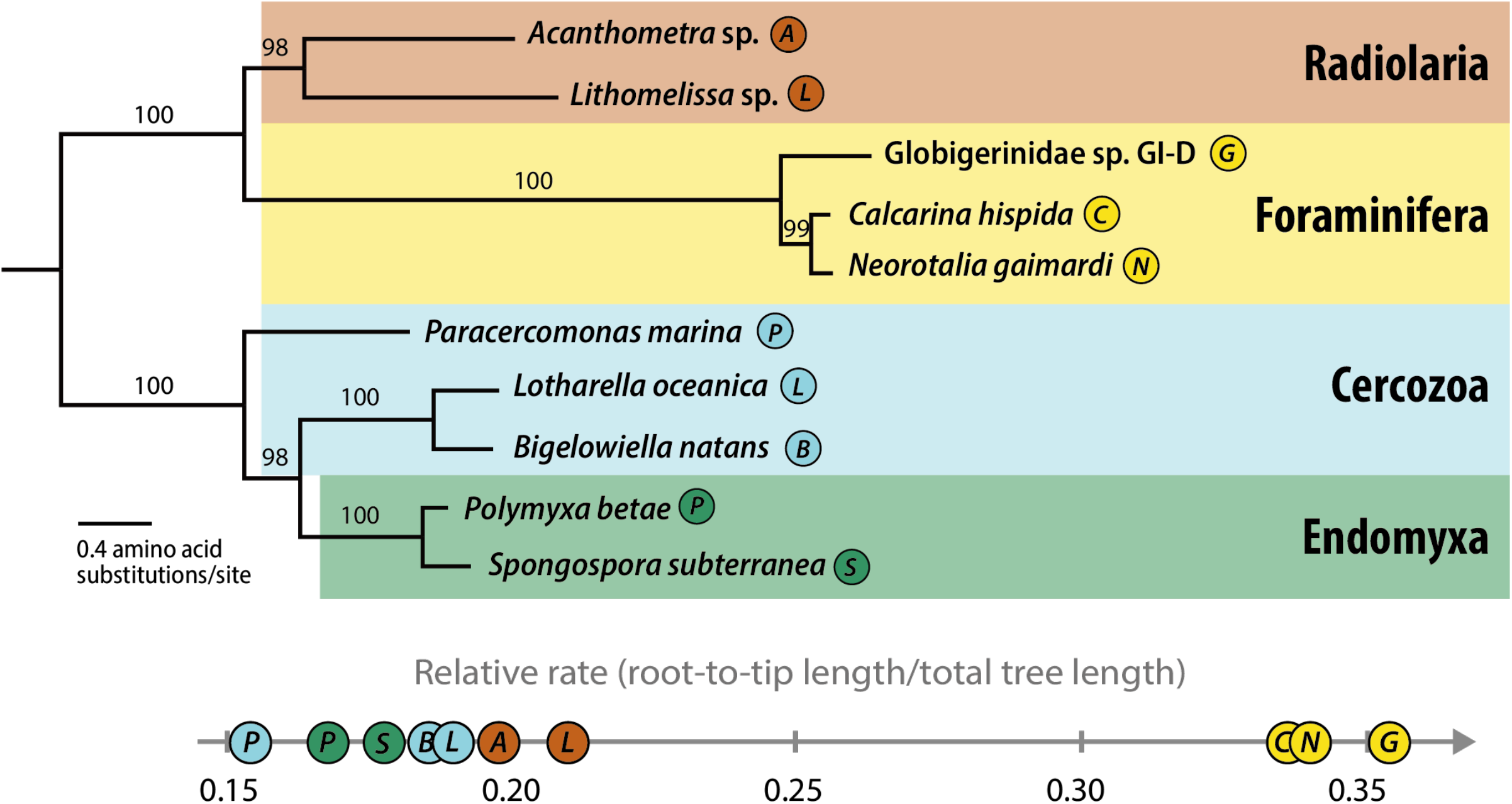
Whole-mitogenome phylogeny of Rhizaria and their relative evolutionary rates. The maximum likelihood tree was constructed using protein sequences predicted from whole mitogenomes and the mtInv+F+R4 model implemented in IQ-TREE. Bootstrap support values (%, based on 1000 replicates) are shown for each branch. Relative evolutionary rates were calculated as the ratio of root-to-tip length to the total tree length, reflecting the proportion of evolutionary change that each branch leads to the overall divergence among sequences. Rooting position is based on Cavalier-Smith et al. [59].

**Figure 4.**
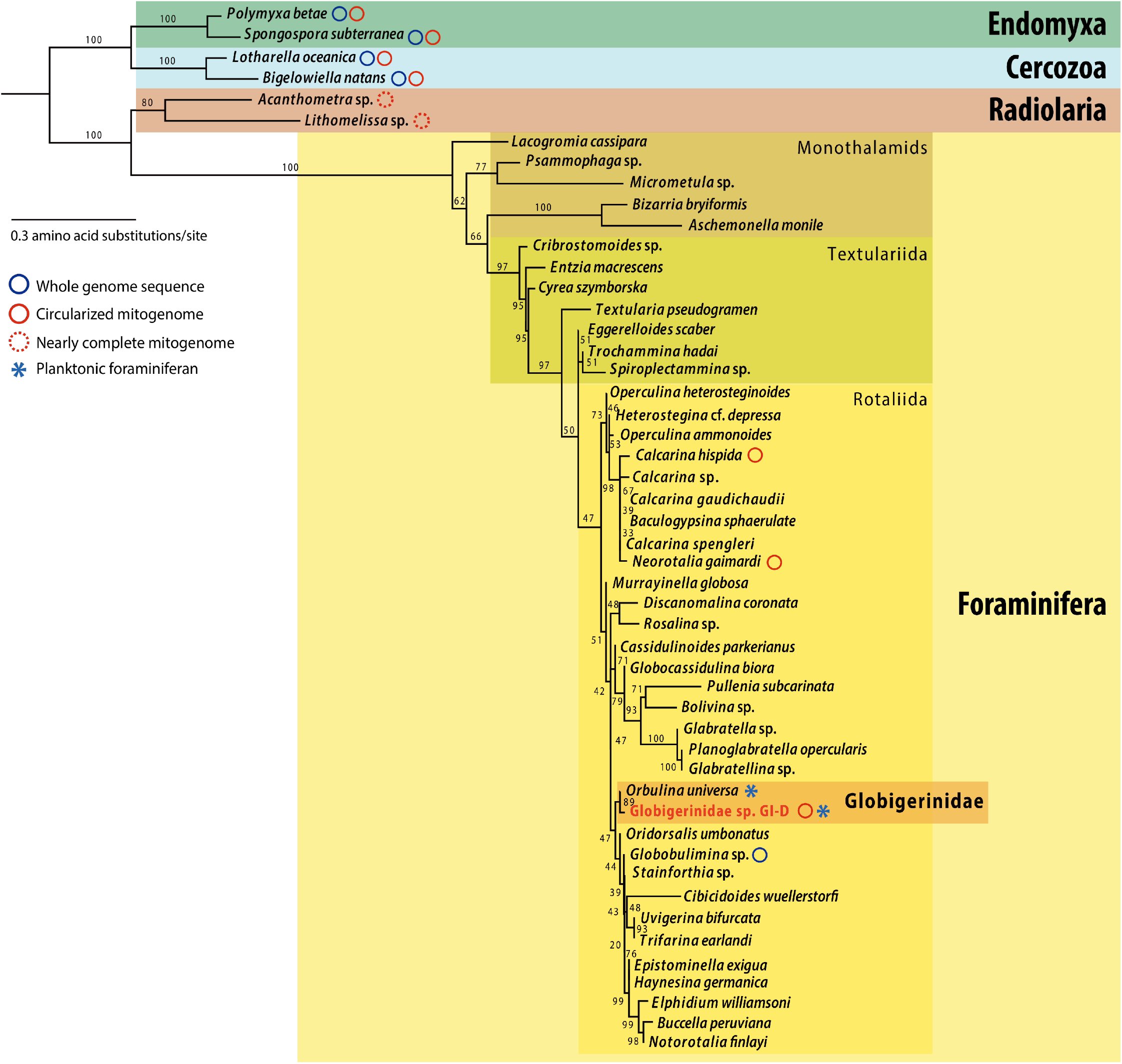
Mitochondrial COX1 phylogeny among Rhizaria. The maximum likelihood tree was reconstructed from COX1 protein sequences of 50 rhizarians (1240 aligned amino acid sites) using IQ-TREE, with the JTT+F+G4 model. Bootstrap support values (%, based on 1000 replicates) are shown for each branch. The planktonic foraminiferan GI-D is the most closely related to *Orbulina universa*, which belongs to the family Globigerinidae (order Rotaliida). Blue circles denote species with whole-genome sequence records, red circles represent the assembly of circularized mitogenomes, and dotted circles indicate the assembly of nearly complete mitogenomes. The planktonic foraminiferans are labeled with asterisks.

Branch length analyses of the phylogenomic tree suggest that the evolutionary rates of foraminiferan mitogenomes are almost twice as fast as those of cercozoans and radiolarians (Figure 3). Further inspection of individual mitochondrial genes reveals that *cox1, cox2, cox3, cob, nad1, nad3, nad4*, nad4L and *nad7* exhibit particularly high evolutionary rates in Foraminifera, while *nad2* shows comparable rates to those observed in cercozoans and radiolarians (**Supplementary Figure 3**).

## Discussion

Through phylogenetic analyses of rhizarian mitogenomes, this study highlights the reductive and accelerated evolution across foraminiferan mitogenomes. Compared with the two previously reported benthic species mitogenomes [25], the mitogenome of the plankton Globigerinidae sp. GI-D has an even smaller size, with greater reduction of protein-coding genes and a lower GC content. Such association between accelerated evolutionary rates, genome reduction, and AT richness is well known in prokaryotic genome evolution [60-61]. The mitogenomes of retarians have completely lost key components, including ribosomal proteins, multiple tRNAs, and ETC genes such as *nad6* and *nad8*. In addition, their ribosomal RNA genes are notably fragmented (**Figure 5**). We note that there is an overall evolutionary trend of gene reduction and accelerated evolution in Rhizaria mitogenomes, from Cercozoa and Endomyxa to Retaria, from Radiolaria to Foramnifera, and from benthic to planktonic foraminiferans (**Figure 6**). The most extreme case of mitogenome reduction is thus represented by the planktonic foraminiferan GI-D, which retains only genes encoding a few components of ETC complexes I and IV.

**Figure 5.**
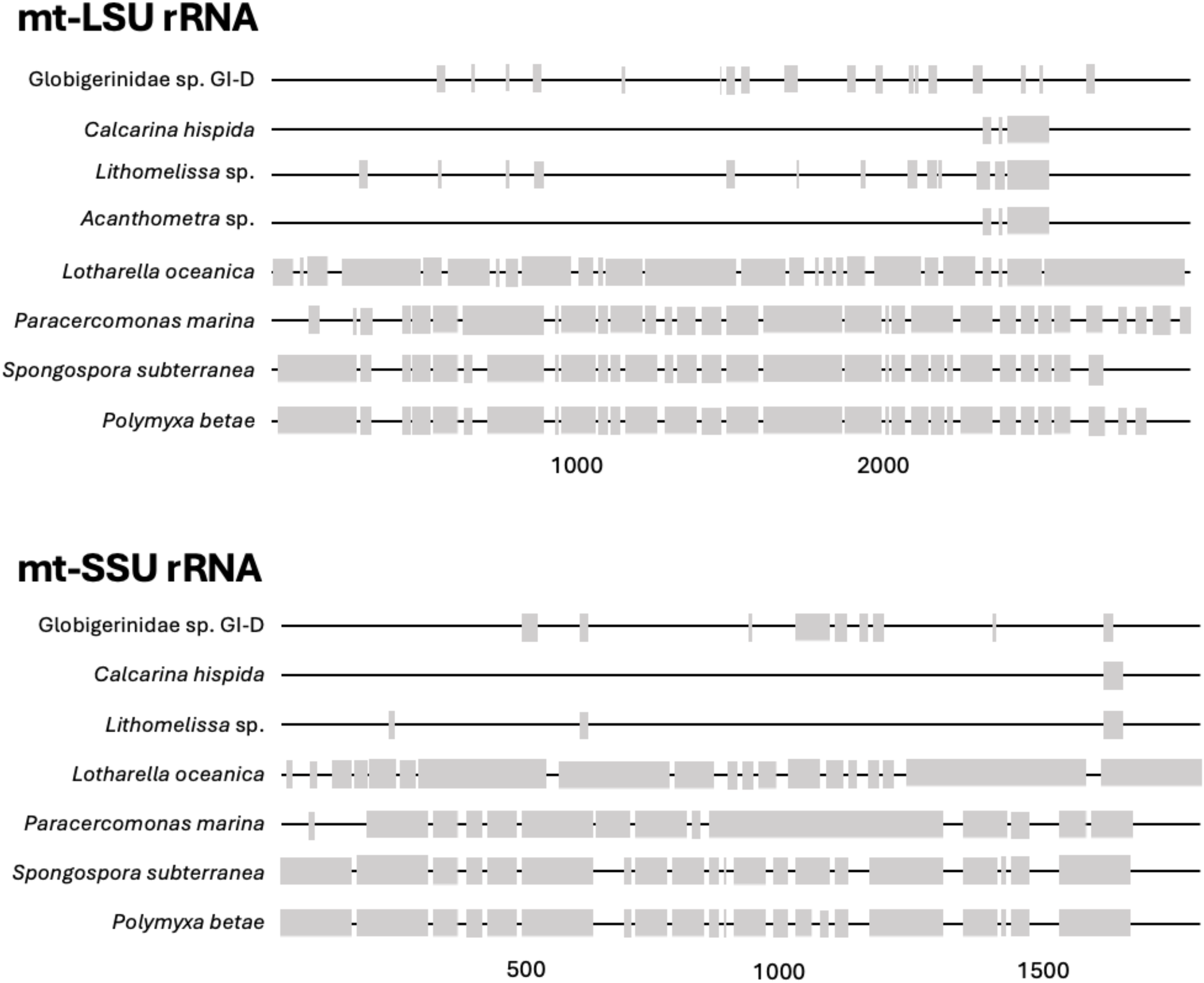
Schematic of mitochondrial ribosomal RNA alignment. Mitochondrial large and small subunit ribosomal RNA genes from representative rhizarians were aligned. Gray blocks represent regions aligned positions, while the black lines indicate unaligned or missing regions. The alignment length (site in bp) is shown along the x-axis.

**Figure 6.**
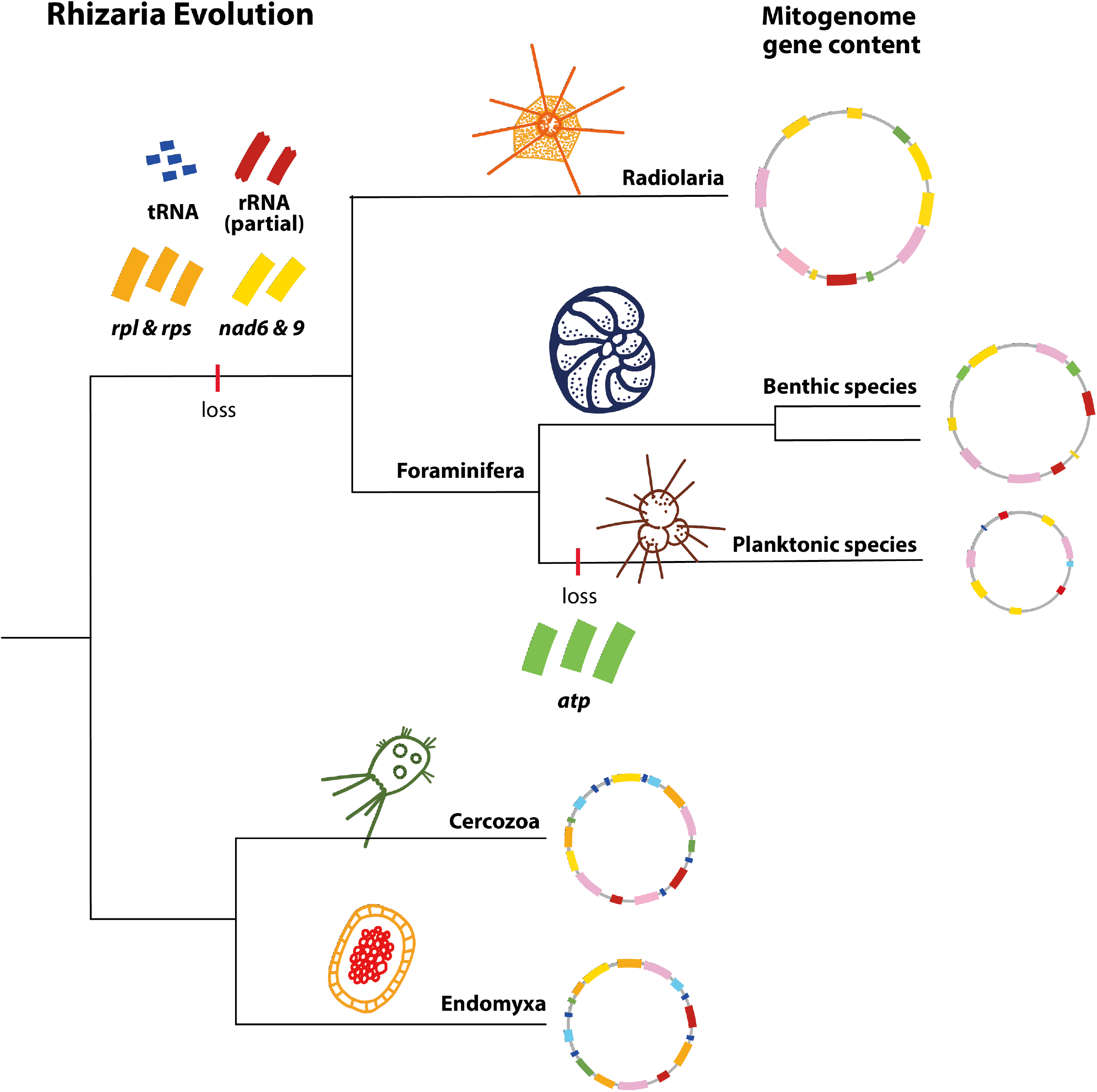
Summary of Rhizaria mitogenome evolution. The mitogenomes of Retaria (Foraminifera & Radiolaria) show a reduction in multiple gene categories, including tRNAs, ribosomal proteins, *nad6, nad9*, and ribosomal RNAs. In planktonic foraminifera, this reduction is even more pronounced, with the complete loss of *atp* subunit genes.

Rapid evolutionary rates have been reported for the nuclear SSU rDNA of planktonic foraminiferans, with gene lengths ranging from 977 bp in *O. universa* to 1178 bp in *Globorotalia menardii* [62]. This accelerated and reductive evolution has been attributed to their shorter reproductive cycles and planktonic lifestyle. In addition, increased molecular evolutionary rates in planktonic microeukaryotes have been linked to fluctuating seawater temperatures, as observed in the genome of marine phytoplankton *Thalassiosira pseudonana* [63]. Given that planktonic foraminiferan inhabit a dynamic ecosystem characterized by variable temperature, salinity and nutrient conditions, their higher sensitivity to environmental fluctuations, including ocean currents and weather patterns [64-65], may contribute to an elevated substitution rate and lineage-specific evolutionary patterns at the molecular level, which is a topic that merits further investigation.

Similar to Foraminifera mitogenomes, pervasive gene loss is observed in another SAR lineage, Myzozoa (Alveolata), which includes apicoplexans, dinoflagellates, and chrompodellids. Myzozoans possess highly reduced mitogenomes, with only ETC complex I and III genes and fragmented ribosomal RNA genes [13,66-67]. This pattern of rRNA fragmentation is also found in distantly related lineages, such as the green alga *Polytomella magna* (Chlorophyta), where a reduced mitochondrial rRNA consists of 13 fragments [68]. A cryo-EM study found that the rRNA in the *Toxoplasma* (apicomplexan) ribosome has over 50 molecules, with repetitive use of the same sequences, addition of poly-A tails, and interaction with non-ribosomal proteins repurposed as components of the mitochondrial ribosome [15]. The widespread convergence of mitochondrial rRNA gene fragmentation and reduction across distant lineages suggests this is a repeated pattern in eukaryote evolution. While myzozoan rRNA fragmentation and massive gene loss have been attributed to an ancestral linearized alveolate mitogenome [14], this may not apply to Foraminifera, given that all currently sequenced Rhizarian mitogenomes are circular. This highlights Foraminifera as an important example for mitogenome evolution. It would be intriguing to investigate if certain cellular or life cycle features are associated with mitogenome reduction and rRNA fragmentation.

Finally, we note that compared with benthic foraminiferans, the GI-D mitogenome has a higher AT content, a higher proportion of repetitive sequences (**Table 2**), and an additional GIY-YIG endonuclease gene (**Figure 1**). Given that repetitive sequences can act as recombination hotspots [69], a higher density of repetitive elements suggests an enhanced potential for recombination-driven genome rearrangement in planktonic foraminiferan mitogenomes. GIY-YIG endonuclease family genes are prevalent in bacteria and have also been identified in the mitogenomes of various eukaryotes, including fungi [70-71] and protists [72]. Given that endonuclease genes have been implicated in DNA cleavage, repair, and recombination [73-74], their presence can also contribute to mitogenome structural changes. This is consistent with the observation that GI-D and the previously reported benthic foraminiferan mitogenomes [25] have little conservation in genome organization. Such mitogenome features and plasticity may be associated with adaptation to environmental stress in marine planktonic environments that are characterized by fluctuations in temperature, oxygen, and other parameters. Overall, our study suggests that planktonic foraminiferans likely represent the pinnacle of mitogenome reductive evolution in Rhizaria, with a small gene repertoire, accelerated evolutionary rates, rRNA fragmentation, low GC content, high repeat density, and low conservation in genome organization. The physiological significance of the reduction and variation of mitogenomes in foraminiferans and the implications of these genomic features for the biology of foraminiferans merit further studies.

## Acknowledgments

We would like to express our sincere gratitude to the Green Island Marine Research Station, Dr. Ming-Jay Ho, Dr. Sophie Nuber, and Dr. Oscar Branson for supporting the field sampling conducted on Green Island. We also thank the Bioinformatics Core Lab, Comprehensive Flow Cytometry Core Lab, Electron Microscopy Division of the Cell Biology Core Lab, and Genomic Technology Core at the Institute of Plant and Microbial Biology, Academia Sinica, Taiwan for their technical support and assistance. This work was supported by an Academia Sinica Career Development Award grant AS-CDA-110-L01 (C.K.) and National Science and Technology Council, Taiwan grants 111-2611-M-001-008-MY3 and 114-2628-B-001-012 (C.K.).

## Supplementary Tables and Figures

**Supplementary Table 1.** Complete or nearly complete rhizarian mitogenome sequences analyzed in this study.

| Species | Accession ID/References | Higher-level taxa | Note |
| --- | --- | --- | --- |
| *Calcarina hispida* | OP965950.1 | Foraminifera |  |
| *Neorotalia gaimardi* | OP965949.1 | Foraminifera |  |
| *Acanthometra* sp. | Macher, J.-N et al., 2023 | Radiolaria | Nearly complete |
| *Lithomelissa* sp. | Macher, J.-N et al., 2023 | Radiolaria | Nearly complete |
| *Polymyxa betae* | NC_059071.1 | Endomyxa |  |
| *Spongospora subterranea* | NC_034004.1 | Endomyxa |  |
| *Lotharella oceanica* | NC_029731.1 | Cercozoa |  |
| *Paracercomonas marina* | KP165385 | Cercozoa |  |
| *Bigelowiella natans* | HQ840955 | Cercozoa | Nearly complete |

**Supplementary Table 2.** Taxonomic distribution of contigs (%) in each planktonic foraminiferan (Globigerinidae) particle SAG

| Particle | GI-A | GI-B | GI-C | GI-D |
| --- | --- | --- | --- | --- |
| Prokaryotic contigs | 97.0% | 84.7% | 74.6% | 68.9% |
| Eukaryotic contigs | 2.7% | 15.1% | 22.5% | 28.9% |
| Unclassified contigs | 0.3% | 0.2% | 2.9% | 2.2% |

**Supplementary Figure 1.**
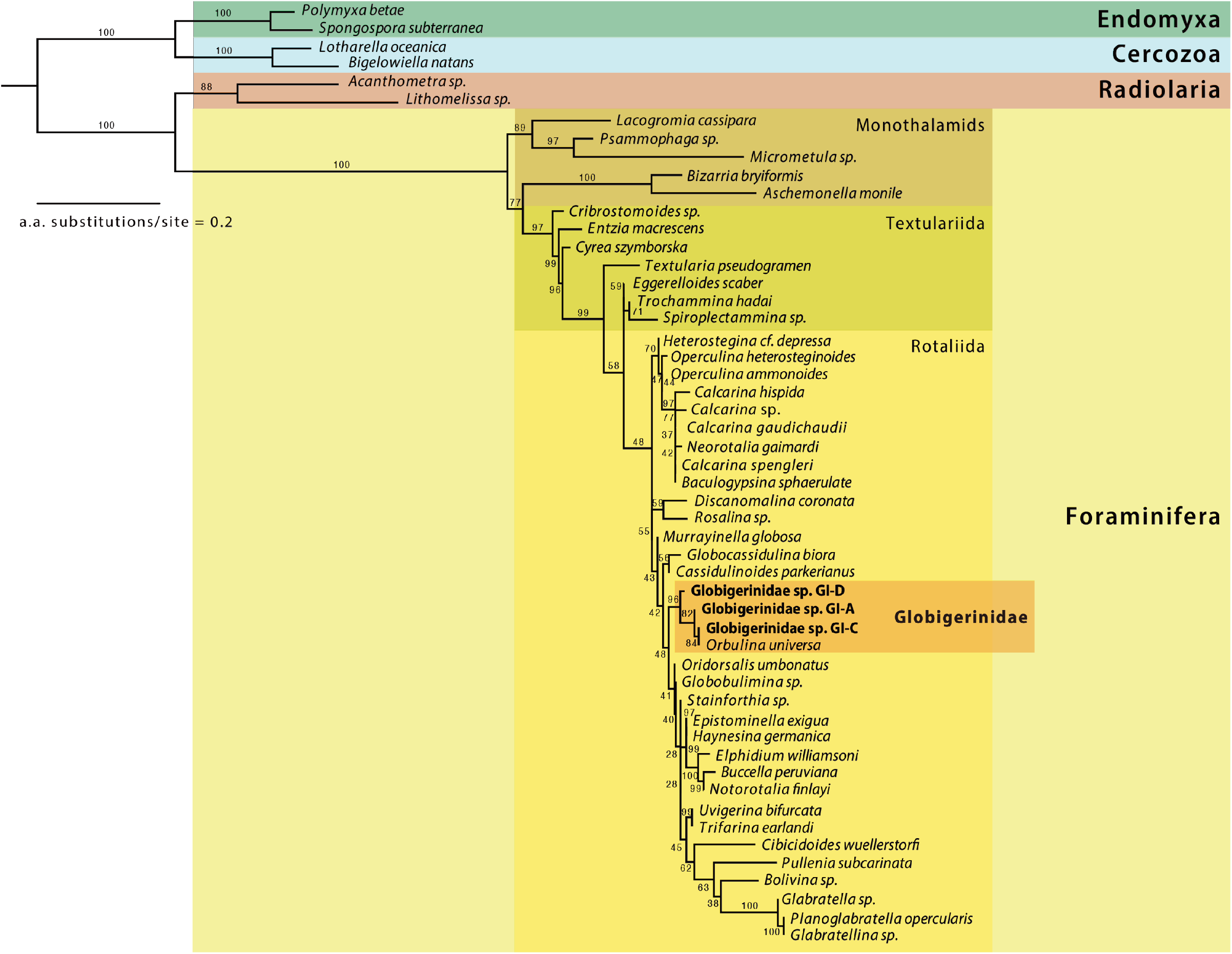
Mitochondrial COX1 protein phylogeny of Foraminifera, including three particles from Green Island (GI) seawater. The maximum likelihood tree was reconstructed from COX1 protein sequences of 52 rhizarians (1240 aligned amino acid sites) using IQ-TREE, with the mtZOA model. Planktonic foraminiferans GI-A, GI-C, and GI-D are clustered with *Orbulina universa* (Globigerinidae, Rotaliida). Bootstrap support values (%, based on 1000 replicates) are shown at the branches.

**Supplementary Figure 2.**
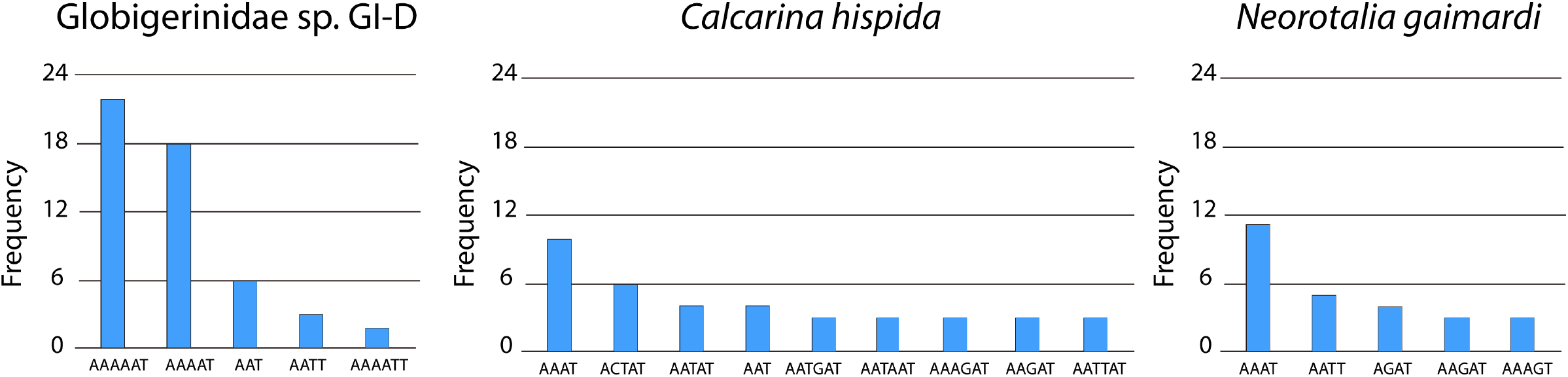
The frequency of repetitive sequences in the mitogenomes of Foraminifera. The bar chart shows the top five frequent repetitive DNA sequences in Globigerinidae sp. GI-D, *C. hispida*, and *N. gaimardi*.

**Supplementary Figure 3.**
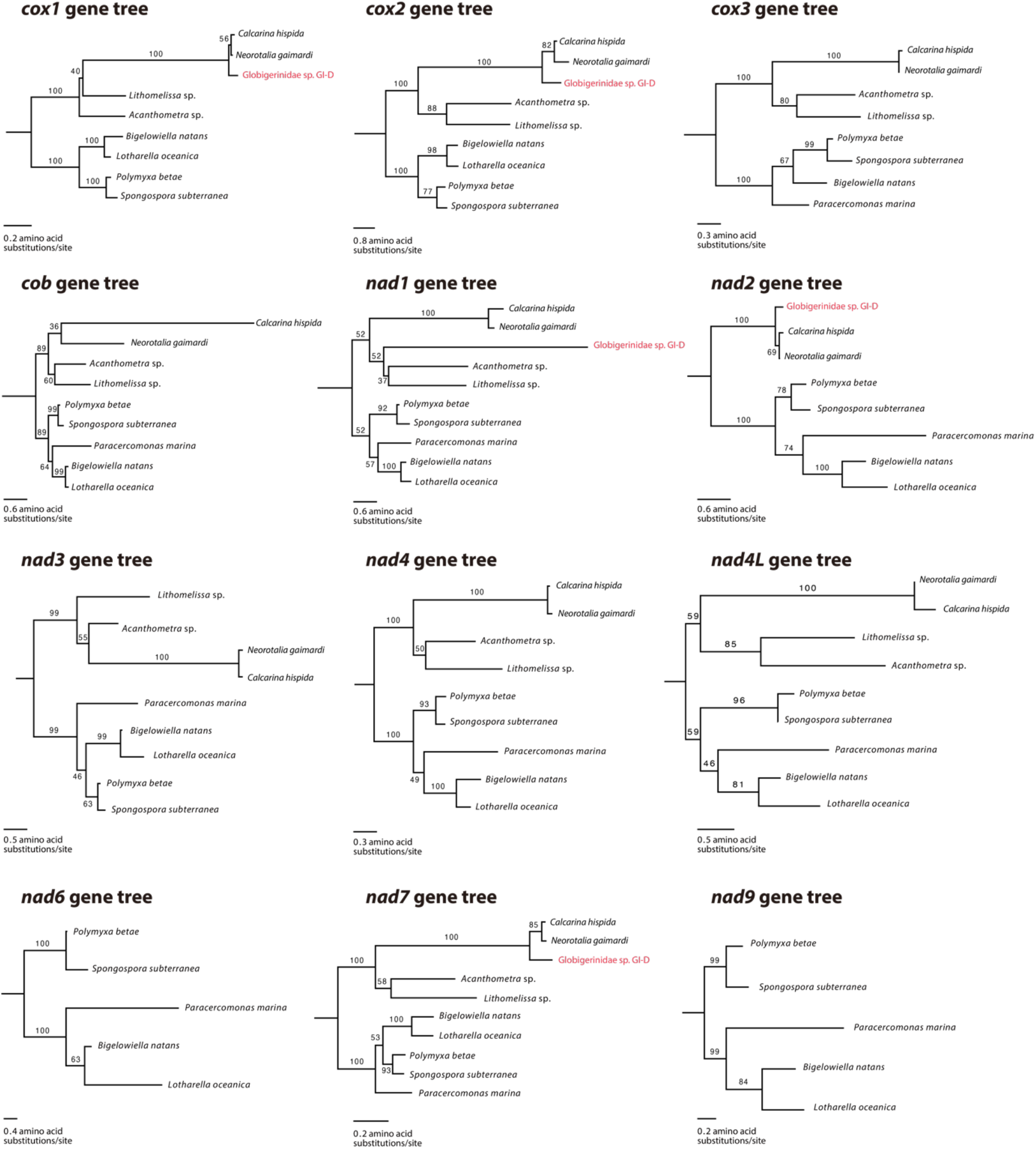
Individual mitochondrial gene trees of Rhizaria. Mitochondrial *cox1, cox2, cob, nad1, nad3, nad4, nad4L*, and *nad7* gene trees reveal an accelerated evolutionary rate in Foraminifera. The trees were constructed based on protein sequences, and the bootstrap support values (%, based on 1000 replicates) are shown at the branches.

## References

1. Zimorski, V., Ku, C., Martin, W. F., & Gould, S. B. (2014). Endosymbiotic theory for organelle origins. Current Opinion in Microbiology, 22, 38–48. 10.1016/j.mib.2014.09.008

2. Archibald, J. M. (2015). Endosymbiosis and Eukaryotic Cell Evolution. Current Biology, 25(19), R911–R921. 10.1016/j.cub.2015.07.055

3. Timmis, J. N., Ayliffe, M. A., Huang, C. Y., & Martin, W. (2004). Endosymbiotic gene transfer: Organelle genomes forge eukaryotic chromosomes. Nature Reviews Genetics, 5(2), 123–135. 10.1038/nrg1271

4. Ku, C., Nelson-Sathi, S., Roettger, M., Sousa, F. L., Lockhart, P. J., Bryant, D., Hazkani-Covo, E., McInerney, J. O., Landan, G., & Martin, W. F. (2015). Endosymbiotic origin and differential loss of eukaryotic genes. Nature, 524(7566), 427–432. 10.1038/nature14963

5. Janouškovec, J., Tikhonenkov, D. V., Burki, F., Howe, A. T., Rohwer, F. L., Mylnikov, A. P., & Keeling, P. J. (2017). A New Lineage of Eukaryotes Illuminates Early Mitochondrial Genome Reduction. Current Biology, 27(23), 3717-3724.e5. 10.1016/j.cub.2017.10.051

6. Butenko, A., Lukeš, J., Speijer, D., & Wideman, J. G. (2024). Mitochondrial genomes revisited: Why do different lineages retain different genes? BMC Biology, 22(1), 15. 10.1186/s12915-024-01824-1

7. Zardoya, R. (2020). Recent advances in understanding mitochondrial genome diversity. F1000Research, 9, 270. 10.12688/f1000research.21490.1

8. Smith, D. R., & Keeling, P. J. (2015). Mitochondrial and plastid genome architecture: Reoccurring themes, but significant differences at the extremes. Proceedings of the National Academy of Sciences, 112(33), 10177–10184. 10.1073/pnas.1422049112

9. Gray, M. W. (2014). The Pre-Endosymbiont Hypothesis: A New Perspective on the Origin and Evolution of Mitochondria. Cold Spring Harbor Perspectives in Biology, 6(3), a016097– a016097. 10.1101/cshperspect.a016097

10. Burger, G., Gray, M. W., Forget, L., & Lang, B. F. (2013). Strikingly Bacteria-Like and Gene-Rich Mitochondrial Genomes throughout Jakobid Protists. Genome Biology and Evolution, 5(2), 418–438. 10.1093/gbe/evt008

11. Müller, M., Mentel, M., Van Hellemond, J. J., Henze, K., Woehle, C., Gould, S. B., Yu, R.-Y., Van Der Giezen, M., Tielens, A. G. M., & Martin, W. F. (2012). Biochemistry and Evolution of Anaerobic Energy Metabolism in Eukaryotes. Microbiology and Molecular Biology Reviews, 76(2), 444–495. 10.1128/MMBR.05024-11

12. Dobáková, E., Flegontov, P., Skalický, T., & Lukeš, J. (2015). Unexpectedly Streamlined Mitochondrial Genome of the Euglenozoan Euglena gracilis. Genome Biology and Evolution, 7(12), 3358–3367. 10.1093/gbe/evv229

13. Waller, R. F., & Jackson, C. J. (2009). Dinoflagellate mitochondrial genomes: Stretching the rules of molecular biology. BioEssays, 31(2), 237–245. 10.1002/bies.200800164

14. Flegontov, P., Michálek, J., Janouškovec, J., Lai, D.-H., Jirků, M., Hajdušková, E., Tomčala, A., Oo, T. D., Keeling, P. J., Pain, A., Oborník, M., & Lukeš, J. (2015). Divergent Mitochondrial Respiratory Chains in Phototrophic Relatives of Apicomplexan Parasites. Molecular Biology and Evolution, 32(5), 1115–1131. 10.1093/molbev/msv021

15. Shikha, S., Tobiasson, V., Ferreira Silva, M., Ovciarikova, J., Beraldi, D., Mühleip, A., & Sheiner, L. (2025). Numerous rRNA molecules form the apicomplexan mitoribosome via repurposed protein and RNA elements. Nature Communications, 16(1), 817. 10.1038/s41467-025-56057-9

16. Adl, S. M., Bass, D., Lane, C. E., Lukeš, J., Schoch, C. L., Smirnov, A., Agatha, S., Berney, C., Brown, M. W., Burki, F., Cárdenas, P., Čepička, I., Chistyakova, L., Del Campo, J., Dunthorn, M., Edvardsen, B., Eglit, Y., Guillou, L., Hampl, V., … Zhang, Q. (2019). Revisions to the Classification, Nomenclature, and Diversity of Eukaryotes. Journal of Eukaryotic Microbiology, 66(1), 4–119. 10.1111/jeu.12691

17. Kucera, M. (2007). Chapter Six Planktonic Foraminifera as Tracers of Past Oceanic Environments. In Developments in Marine Geology (Vol. 1, pp. 213–262). Elsevier. 10.1016/S1572-5480(07)01011-1

18. Yasuhara, M., Tittensor, D. P., Hillebrand, H., & Worm, B. (2017). Combining marine macroecology and palaeoecology in understanding biodiversity: Microfossils as a model: Marine macroecology-palaeoecology integration. Biological Reviews, 92(1), 199–215. 10.1111/brv.12223

19. Pawlowski, J., Holzmann, M., & Tyszka, J. (2013). New supraordinal classification of Foraminifera: Molecules meet morphology. Marine Micropaleontology, 100, 1–10. 10.1016/j.marmicro.2013.04.002

20. Morard, R., Darling, K. F., Weiner, A. K. M., Hassenrück, C., Vanni, C., Cordier, T., Henry, N., Greco, M., Vollmar, N. M., Milivojevic, T., Rahman, S. N., Siccha, M., Meilland, J., Jonkers, L., Quillévéré, F., Escarguel, G., Douady, C. J., De Garidel-Thoron, T., De Vargas, C., & Kucera, M. (2024). The global genetic diversity of planktonic foraminifera reveals the structure of cryptic speciation in plankton. Biological Reviews, 99(4), 1218–1241. 10.1111/brv.13065

21. Macher, J.-N., Bloska, D. M., Holzmann, M., Girard, E. B., Pawlowski, J., & Renema, W. (2022). Mitochondrial cytochrome c oxidase subunit I (COI) metabarcoding of Foraminifera communities using taxon-specific primers. PeerJ, 10, e13952. 10.7717/peerj.13952

22. Habura, A., Hou, Y., Reilly, A. A., & Bowser, S. S. (2011). High-throughput sequencing of Astrammina rara: Sampling the giant genome of a giant foraminiferan protist. BMC Genomics, 12(1), 169. 10.1186/1471-2164-12-169

23. Woehle, C., Roy, A.-S., Glock, N., Wein, T., Weissenbach, J., Rosenstiel, P., Hiebenthal, C., Michels, J., Schönfeld, J., & Dagan, T. (2018). A Novel Eukaryotic Denitrification Pathway in Foraminifera. Current Biology, 28(16), 2536-2543.e5. 10.1016/j.cub.2018.06.027

24. Glöckner, G., Hülsmann, N., Schleicher, M., Noegel, A. A., Eichinger, L., Gallinger, C., Pawlowski, J., Sierra, R., Euteneuer, U., Pillet, L., Moustafa, A., Platzer, M., Groth, M., Szafranski, K., & Schliwa, M. (2014). The Genome of the Foraminiferan Reticulomyxa filosa. Current Biology, 24(1), 11–18. 10.1016/j.cub.2013.11.027

25. Macher, J.-N., Coots, N. L., Poh, Y.-P., Girard, E. B., Langerak, A., Muñoz-Gómez, S. A., Sinha, S. D., Jirsová, D., Vos, R., Wissels, R., Gile, G. H., Renema, W., & Wideman, J. G. (2023). Single-Cell Genomics Reveals the Divergent Mitochondrial Genomes of Retaria (Foraminifera and Radiolaria). mBio, 14(2), e00302–23. 10.1128/mbio.00302-23

26. Ku, C., & Sebé-Pedrós, A. (2019). Using single-cell transcriptomics to understand functional states and interactions in microbial eukaryotes. Philosophical Transactions of the Royal Society B: Biological Sciences, 374(1786), 20190098. 10.1098/rstb.2019.0098

27. Ciobanu, D., Clum, A., Ahrendt, S., Andreopoulos, W. B., Salamov, A., Chan, S., Quandt, C. A., Foster, B., Meier-Kolthoff, J. P., Tang, Y. T., Schwientek, P., Benny, G. L., Smith, M. E., Bauer, D., Deshpande, S., Barry, K., Copeland, A., Singer, S. W., Woyke, T., … Cheng, J.-F. (2021). A single-cell genomics pipeline for environmental microbial eukaryotes. iScience, 24(4), 102290. 10.1016/j.isci.2021.102290

28. Keeling, P. J. (2019). Combining morphology, behaviour and genomics to understand the evolution and ecology of microbial eukaryotes. Philosophical Transactions of the Royal Society B: Biological Sciences, 374(1786), 20190085. 10.1098/rstb.2019.0085

29. Schön, M. E., Zlatogursky, V. V., Singh, R. P., Poirier, C., Wilken, S., Mathur, V., Strassert, J. F. H., Pinhassi, J., Worden, A. Z., Keeling, P. J., Ettema, T. J. G., Wideman, J. G., & Burki, F. (2021). Single cell genomics reveals plastid-lacking Picozoa are close relatives of red algae. Nature Communications, 12(1), 6651. 10.1038/s41467-021-26918-0

30. Wideman, J. G., Monier, A., Rodríguez-Martínez, R., Leonard, G., Cook, E., Poirier, C., Maguire, F., Milner, D. S., Irwin, N. A. T., Moore, K., Santoro, A. E., Keeling, P. J., Worden, A. Z., & Richards, T. A. (2019). Unexpected mitochondrial genome diversity revealed by targeted single-cell genomics of heterotrophic flagellated protists. Nature Microbiology, 5(1), 154– 165. 10.1038/s41564-019-0605-4

31. Záhonová, K., Lax, G., Sinha, S. D., Leonard, G., Richards, T. A., Lukeš, J., & Wideman, J. G. (2021). Single-cell genomics unveils a canonical origin of the diverse mitochondrial genomes of euglenozoans. BMC Biology, 19(1), 103. 10.1186/s12915-021-01035-y

32. Mohamed Yusoff, A. A., Mohd Khair, S. Z. N., & Abd Radzak, S. M. (2025). Mitochondrial DNA copy number alterations: Key players in the complexity of glioblastoma (Review). Molecular Medicine Reports, 31(3), 78. 10.3892/mmr.2025.13443

33. Cai-Li, R.-Y., Ren, H., Fang, W.-N., Yang, E.-W., Chen, W.-H., LeKieffre, C., Branson, O., Fehrenbacher, J., Ve_er, L., Jeng, M.-S., & Spero, H. J. (2025). Symbiont regulation of nitrogen metabolism and excretion in tropical planktonic foraminifera. Geochimica et Cosmochimica Acta, S0016703725001255. 10.1016/j.gca.2025.03.009

34. Fang, W.-N., Branson, O., Yang, E.-W., Chen, W.-H., Cai-Li, R.-Y., Spero, H. J., Fehrenbacher, J., Ve_er, L., LeKieffre, C., & Ren, H. (2025). Direct pathway of incorporating dietary nitrogen in shell-bound matrix of the planktic foraminifera Trilobatus sacculifer. Earth and Planetary Science LeUers, 654, 119231. 10.1016/j.epsl.2025.119231

35. Chen, S., Zhou, Y., Chen, Y., & Gu, J. (2018). fastp: An ultra-fast all-in-one FASTQ preprocessor. Bioinformatics, 34(17), i884–i890. 10.1093/bioinformatics/bty560

36. Wick, R. R., Judd, L. M., & Holt, K. E. (2018). Deepbinner: Demultiplexing barcoded Oxford Nanopore reads with deep convolutional neural networks. PLOS Computational Biology, 14(11), e1006583. 10.1371/journal.pcbi.1006583

37. Prjibelski, A., Antipov, D., Meleshko, D., Lapidus, A., & Korobeynikov, A. (2020). Using SPAdes De Novo Assembler. Current Protocols in Bioinformatics, 70(1), e102. 10.1002/cpbi.102

38. Karlicki, M., Antonowicz, S., & Karnkowska, A. (2022). Tiara: Deep learning-based classification system for eukaryotic sequences. Bioinformatics, 38(2), 344–350. 10.1093/bioinformatics/btab672

39. Camacho, C., Coulouris, G., Avagyan, V., Ma, N., Papadopoulos, J., Bealer, K., & Madden, T. L. (2009). BLAST+: Architecture and applications. BMC Bioinformatics, 10(1), 421. 10.1186/1471-2105-10-421

40. Tanifuji, G., Archibald, J. M., & Hashimoto, T. (2016). Comparative genomics of mitochondria in chlorarachniophyte algae: Endosymbiotic gene transfer and organellar genome dynamics. Scientific Reports, 6(1), 21016. 10.1038/srep21016

41. Dierckxsens, N., Mardulyn, P., & Smits, G. (2016). NOVOPlasty: De novo assembly of organelle genomes from whole genome data. Nucleic Acids Research, gkw955. 10.1093/nar/gkw955

42. Langmead, B., & Salzberg, S. L. (2012). Fast gapped-read alignment with Bowtie 2. Nature Methods, 9(4), 357–359. 10.1038/nmeth.1923

43. Li, H. (2018). Minimap2: Pairwise alignment for nucleotide sequences. Bioinformatics, 34(18), 3094–3100. 10.1093/bioinformatics/bty191

44. Thorvaldsdottir, H., Robinson, J. T., & Mesirov, J. P. (2013). Integrative Genomics Viewer (IGV): High-performance genomics data visualization and exploration. Briefings in Bioinformatics, 14(2), 178–192. 10.1093/bib/bbs017

45. Gurevich, A., Saveliev, V., Vyahhi, N., & Tesler, G. (2013). QUAST: Quality assessment tool for genome assemblies. Bioinformatics, 29(8), 1072–1075. 10.1093/bioinformatics/btt086

46. Avvaru, A. K., Sowpati, D. T., & Mishra, R. K. (2018). PERF: An exhaustive algorithm for ultra-fast and efficient identification of microsatellites from large DNA sequences. Bioinformatics, 34(6), 943–948. 10.1093/bioinformatics/btx721

47. Lang, B. F., Beck, N., Prince, S., Sarrasin, M., Rioux, P., & Burger, G. (2023). Mitochondrial genome annotation with MFannot: A critical analysis of gene identification and gene model prediction. Frontiers in Plant Science, 14, 1222186. 10.3389/fpls.2023.1222186

48. Bernt, M., Donath, A., Jühling, F., Externbrink, F., Florentz, C., Fritzsch, G., Pütz, J., Middendorf, M., & Stadler, P. F. (2013). MITOS: Improved de novo metazoan mitochondrial genome annotation. Molecular Phylogenetics and Evolution, 69(2), 313–319. 10.1016/j.ympev.2012.08.023

49. Greiner, S., Lehwark, P., & Bock, R. (2019). OrganellarGenomeDRAW (OGDRAW) version 1.3.1: Expanded toolkit for the graphical visualization of organellar genomes. Nucleic Acids Research, 47(W1), W59–W64. 10.1093/nar/gkz238

50. Emms, D. M., & Kelly, S. (2019). OrthoFinder: Phylogenetic orthology inference for comparative genomics. Genome Biology, 20(1), 238. 10.1186/s13059-019-1832-y

51. Nguyen, L.-T., Schmidt, H. A., Von Haeseler, A., & Minh, B. Q. (2015). IQ-TREE: A Fast and Effective Stochastic Algorithm for Estimating Maximum-Likelihood Phylogenies. Molecular Biology and Evolution, 32(1), 268–274. 10.1093/molbev/msu300

52. Kalyaanamoorthy, S., Minh, B. Q., Wong, T. K. F., Von Haeseler, A., & Jermiin, L. S. (2017). ModelFinder: Fast model selection for accurate phylogenetic estimates. Nature Methods, 14(6), 587–589. 10.1038/nmeth.4285

53. Katoh, K., & Standley, D. M. (2013). MAFFT Multiple Sequence Alignment Software Version 7: Improvements in Performance and Usability. Molecular Biology and Evolution, 30(4), 772– 780. 10.1093/molbev/mst010

54. Rambaut, A. (2012). FigTree v1. 4. Molecular evolution, phylogenetics andepidemiology. Edinburgh: University of Edinburgh, Institute of Evolutionary Biology.

55. Carpenter, W. B., Parker, W. R., Jones, T. R., & Hardwicke, R. (1862). Introduction to the study of the foraminifera. Journal of Cell Science, S2-2(8), 297–301. 10.1242/jcs.s2-2.8.297

56. Chinnery, P. F., & Hudson, G. (2013). Mitochondrial genetics. British Medical Bulletin, 106(1), 135–159. 10.1093/bmb/ldt017

57. Pentinsaari, M., Salmela, H., Mutanen, M., & Roslin, T. (2016). Molecular evolution of a widely-adopted taxonomic marker (COI) across the animal tree of life. Scientific Reports, 6(1), 35275. 10.1038/srep35275

58. Orbigny, A.D.d’. (1839). Foraminifères [of Cubati (Vol. 8). Bertrand.

59. Cavalier-Smith, T., Chao, E. E., & Lewis, R. (2018). Multigene phylogeny and cell evolution of chromist infrakingdom Rhizaria: Contrasting cell organisation of sister phyla Cercozoa and Retaria. Protoplasma, 255(5), 1517–1574. 10.1007/s00709-018-1241-1

60. Dufresne, A., Garczarek, L., & Partensky, F. (2005). Accelerated evolution associated with genome reduction in a free-living prokaryote. Genome Biology, 6(2), R14. 10.1186/gb-2005-6-2-r14

61. McCutcheon, J. P., & Moran, N. A. (2012). Extreme genome reduction in symbiotic bacteria. Nature Reviews Microbiology, 10(1), 13–26. 10.1038/nrmicro2670

62. de Vargas, C., Zaninetti, L., Hilbrecht, H., & Pawlowski, J. (1997). Phylogeny and Rates of Molecular Evolution of Planktonic Foraminifera: SSU rDNA Sequences Compared to the Fossil Record. Journal of Molecular Evolution, 45(3), 285–294. 10.1007/PL00006232

63. Schaum, C.-E., Buckling, A., Smirnoff, N., Studholme, D. J., & Yvon-Durocher, G. (2018). Environmental fluctuations accelerate molecular evolution of thermal tolerance in a marine diatom. Nature Communications, 9(1), 1719. 10.1038/s41467-018-03906-5

64. Wei, K., & Kennett, J. P. (1986). Taxonomic evolution of Neogene planktonic foraminifera and paleoceanographic relations. Paleoceanography, 1(1), 67–84. 10.1029/PA001i001p00067

65. Malmgren, B. A., & Berggren, W. A. (1987). Evolutionary changes in some Late Neogene planktonic foraminiferal lineages and their relationships to paleoceanographic changes. Paleoceanography, 2(5), 445–456. 10.1029/PA002i005p00445

66. Nash, E. A., Nisbet, R. E. R., Barbrook, A. C., & Howe, C. J. (2008). Dinoflagellates: A mitochondrial genome all at sea. Trends in Genetics, 24(7), 328–335. 10.1016/j.tig.2008.04.001

67. Feagin, J. E., Harrell, M. I., Lee, J. C., Coe, K. J., Sands, B. H., Cannone, J. J., Tami, G., Schnare, M. N., & Gutell, R. R. (2012). The Fragmented Mitochondrial Ribosomal RNAs of Plasmodium falciparum. PLoS ONE, 7(6), e38320. 10.1371/journal.pone.0038320

68. Tobiasson, V., Berzina, I., & Amunts, A. (2022). Structure of a mitochondrial ribosome with fragmented rRNA in complex with membrane-targeting elements. Nature Communications, 13(1), 6132. 10.1038/s41467-022-33582-5

69. Aras, R. A., Kang, J., Tschumi, A. I., Harasaki, Y., & Blaser, M. J. (2003). Extensive repetitive DNA facilitates prokaryotic genome plasticity. Proceedings of the National Academy of Sciences, 100(23), 13579–13584. 10.1073/pnas.1735481100

70. Saguez, C. (2000). Intronic GIY-YIG endonuclease gene in the mitochondrial genome of Podospora curvicolla: Evidence for mobility. Nucleic Acids Research, 28(6), 1299–1306. 10.1093/nar/28.6.1299

71. Megarioti, A. H., & Kouvelis, V. N. (2020). The Coevolution of Fungal Mitochondrial Introns and Their Homing Endonucleases (GIY-YIG and LAGLIDADG). Genome Biology and Evolution, 12(8), 1337–1354. 10.1093/gbe/evaa126

72. Craig, R. J., Yushenova, I. A., Rodriguez, F., & Arkhipova, I. R. (2021). An Ancient Clade of Penelope -Like Retroelements with Permuted Domains Is Present in the Green Lineage and Protists, and Dominates Many Invertebrate Genomes. Molecular Biology and Evolution, 38(11), 5005–5020. 10.1093/molbev/msab225

73. Aravind, L. (1999). Conserved domains in DNA repair proteins and evolution of repair systems. Nucleic Acids Research, 27(5), 1223–1242. 10.1093/nar/27.5.1223

74. Mak, A. N.-S., Lambert, A. R., & Stoddard, B. L. (2010). Folding, DNA Recognition, and Function of GIY-YIG Endonucleases: Crystal Structures of R.Eco29kI. Structure, 18(10), 1321–1331. 10.1016/j.str.2010.07.006

